# Divergent paths, convergent heads: morphological adaptation of head shape to habitat use and diet in snakes

**DOI:** 10.1101/2025.05.17.654544

**Authors:** David Hudry, Anthony Herrel

## Abstract

Morphological convergence—where distantly related species evolve similar traits in response to shared ecological pressures—is a hallmark of adaptive evolution. In snakes, head shape is a key functional trait linked to habitat use and diet, yet the extent and drivers of its convergence across habitat types and diet remain poorly understood. Here, we used geometric morphometric analyses to analyze dorsal head shape from photographs of over 266 snake species, representing ∼5% of global snake diversity. Our analyses reveal that head shape variation is only weakly structured by phylogeny and allometry, but shaped by ecological specialization. Morphospace patterns reflect distinct differences in species with different ecologies: fossorial species exhibit compact, posteriorly wider heads suited for burrowing; aquatic species show streamlined profiles for hydrodynamic efficiency; and arboreal snakes tend to possess elongated heads for maneuvering in complex habitats. Terrestrial and semi-fossorial snakes display broad morphospace overlap and an elevated shape disparity highlighting morphological versatility. While diet covaried significantly with head shape, species with similar diets did not exhibit strong morphological convergence. These findings underscore the dominant role of ecology over phylogeny in shaping the evolution of head shape in gape-limited predators like snakes.

## Introduction

The environment plays a central role in shaping animal morphology by favoring traits that enhance survival and reproductive success. When species are exposed to similar ecological pressures, they often evolve similar phenotypes—an evolutionary pattern known as convergent evolution (Losos, 2011; Stayton, 2015). However, the extent to which traits respond to ecological selection can vary significantly across morphological systems (Wainwright, 2007; Deepak et al., 2022). Different factors can explain head shape variation including feeding behavior, sexual selection, and habitat use (Pough & Groves, 1983; Moon et al., 2019). Species with specialized diets, such as mollusks feeders, may, for example evolve asymmetric mandibles in response to whether snails are left or right wound (Pandelis et al., 2023). Sexual selection may also impact head shape, especially if it impacts mate attraction or competitive ability (Fabre et al., 2016). However, most often a multitude of sometimes conflicting selective pressures drive the evolution of any given trait (Huey & Kingsolver, 1989). Consequently, species inhabiting similar habitats may follow distinct evolutionary and morphological pathways (Losos et al., 1998). This is especially true for aquatic snakes facing similar environmental pressures, yet showing different body plans and head shapes (Fabre et al., 2016; Segall et al., 2020; Sheratt et al., 2018). Moreover, despite similar ecological pressures, head shapes may diverge across different clades because other factors such as evolutionary history and habitat-specific behaviors also shape morphology (Speed & Arbuckle, 2017; Grundler & Rabosky, 2014).

Snakes are an excellent model system to study morphological convergence because of their ecological versatility (Pyron & Burbrink, 2012) and their reliance on the head to feed as gape-limited predators (Moon et al., 2019). Among the various selective pressures acting on snake morphology, diet is by far the most studied and influential driver of head shape evolution. Numerous studies have demonstrated that trophic specialization—whether targeting elongate, bulky, hard-shelled, or elusive prey—has led to distinct and repeated morphological adaptations in cranial morphology (Klaczko et al., 2016; Fabre et al., 2016; Segall et al., 2020; Deepak et al., 2022). For example, feeding-related morphological evolution is evident in sea snakes, where selective pressures related to foraging mode have resulted in distinct head shapes (Sherratt et al., 2018). Similarly, homalopsid and xenodontine snakes have evolved unique head shapes driven by the functional demands imposed by the prey they eat (Klaczko et al., 2016; Fabre et al., 2016). Habitat-related selective pressures also influence snake morphology. *Boidae* and *Pythonidae*, for instance, exhibit head shape convergence driven primarily by habitat use (Esquerré & Keogh, 2016).

Previous studies on head shape convergence in snakes have focused on arboreal, aquatic, fossorial and terrestrial snakes. For instance, aquatic species frequently develop narrow heads with upward-facing eyes and nostrils, features that improve their ability to capture prey, move efficiently through water and spot predators while remaining submerged (Hibbitts and Fitzgerald, 2005; Vincent et al., 2009; Segall et al., 2016, 2020; Deepak et al., 2022). Similarly, burrowing snakes possess robust heads adapted for digging and streamlined bodies with reduced ventral scales, which help them burrow with minimal friction (Fabre et al., 2016; Herrel et al., 2008). Meanwhile, microcephalic sea snakes exhibit distinct head shapes linked to developmental changes, representing habitat specialization rather than direct functional convergence (Sherratt et al., 2019). Terrestrial snakes from different lineages often exhibit convergence in robust, broad head shapes associated with subduing large or ground-dwelling prey, reflecting similar ecological demands despite phylogenetic differences (Klaczko et al., 2016; da Silva et al., 2018; Carrasco et al., 2023; Pandelis et al., 2023).

Finally, arboreal snakes frequently converge on slender, elongate head shapes with enlarged orbits—adaptations that enhance precision in prey capture and navigation in structurally complex habitats (Carrasco et al., 2023; Pandelis et al., 2023; Seneci et al., 2025).

Our overarching prediction is that snakes inhabiting similar environments or eating similar prey will show morphological convergence. Particularly, we predict that fossorial snakes will exhibit a narrow, V-shaped head that facilitates substrate penetration and minimizes the energetic cost of burrowing (Navas et al., 2004). This prediction is supported by recent work showing that fossorial snakes display consistent cranial streamlining associated with head-first burrowing, including reduced head width and reinforced snouts (Deepak et al., 2022; Seneci et al., 2025). These morphological traits have evolved independently in multiple lineages of burrowing snakes, reflecting strong functional convergence related to locomotion in compact substrates. Head shape in fossorial snakes is expected to be primarily shaped by locomotor demands rather than feeding ecology, as these species typically consume small, soft-bodied soil invertebrates that do not require specialized cranial adaptations for prey capture or processing. Arboreal snakes are predicted to have slender heads, which enhance crypsis and maneuverability among branches and foliage— traits commonly observed in arboreal colubrids and vipers (Carrasco et al., 2023; Palci et al., 2024). However, mollusk specialists such as *Dipsas* and *Sibon* often exhibit more robust or blunt head shapes, which are associated with the biomechanical demands of molluscivory, including increased muscle mass and cranial rigidity for extracting soft-bodied prey from shells (Palci et al., 2024). Aquatic snakes are expected to exhibit high disparity in head shape, reflecting their exploitation of diverse prey types and the functional demands of navigating various aquatic environments (Deepak et al., 2022). This pattern has been documented in multiple aquatic snake lineages, where variation in prey capture strategies (e.g., gripping, constriction) and ecological specialization (e.g., fish, amphibians, crustaceans) correlates with significant cranial morphological diversity (Segall et al., 2016). Species that utilize multiple environments (e.g., semi-aquatic, semi-arboreal, semi-fossorial) are predicted to exhibit intermediate morphological features that reflect functional compromises between distinct ecological demands. These semi-specialized species would occupy broader morphospaces due to the need to balance locomotion and foraging strategies across habitats (Ortiz-Medina et al., 2022; Pandelis et al., 2023).We also predict that greater morphological variation will be observed in snakes that live in transitional habitats and that have non-specialized diets.

## Materials and Methods

### Sample and ecological classification

We collected data from 281 species used for habitat use (521 specimens) and 197 species for diet (384 specimens). Animals were housed in three European Natural History Museums: the research collection of the last author (99 specimens), the Alexander Koenig Museum in Bonn (319 specimens), and the Museum of Natural Sciences in Brussels (103 specimens). Our measurements were taken from adult specimens whenever possible.

However, in some cases, only young individuals were available. For some species, only subadults were measured because of a lack of availability of adults (Table S1). Our sampling aimed to encompass the phylogenetic and ecological diversity of snakes, while remaining limited by the availability of specimens in these collections.

Based on the literature, eight habitat use categories were determined (Spawls et al., 2002; Whitaker & Captain, 2004; Campbell & Lamar, 2004; Marais, 2004; O’Shea, 2005; Vogel, 2006; Glaw & Vences, 2007; Bartlett & Bartlett, 2009; Das, 2010; O’Shea, 2018; Allen, 2019; Chippaux, 2019; Pietersen et al., 2021; Egan, 2022; Das, 2023). We classified the species into 102 terrestrial, 34 arboreal, 19 aquatic, 7 fossorial, 43 semi-arboreal, 33 semi-aquatic, and 29 semi-fossorial species. We based our ecological categorization on the biomechanical constraints imposed by the environment (Table S2). For instance, similar hydrodynamic constraints affect both marine and freshwater snakes leading us to group them into the same category.

We used the time-calibrated phylogeny and dietary data from the supplementary material of the paper by Title et al. (2024). This paper contain a recent time-calibrated phylogeny of 6,885 species and a complete dataset of diet proportions of more than 68,547 species of squamates. Categories were modified and regrouped based on the functional constraints imposed by the prey. We define a total of ten dietary groups, including 19 generalists, 3 hard egg feeders, 76 hard prey feeders, 16 invertebrate feeders, 2 mollusk specialists, 2 soft egg specialists, and 79 soft prey eaters (Table S3). In creating these categories, we grouped salamanders, lizards, and fish together as they have cylindrical and flexible bodies and as such should impose similar functional constraints.

### Geometric morphometric data

For each individual a photo of the head in dorsal view was taken with a Nikon © D3100 camera. We omitted photos from a lateral view because of sample quality (head deformed & specimens with mouth partially open). Snakes were placed on a grid paper for calibration and scale (1 x 1cm squares; Figure S1A). To perform the geometric morphometric analysis, 25 landmarks and 10 semi-landmarks were selected (Figure S1B) and defined (Table S4). These landmarks and semi-landmarks were based on the literature (Manier, 2004; Mangiacotti et al., 2014; Ruane, 2015; Roth-Monzón et al., 2021). They capture the general head shape of the animals and are easy to identify. This approach was applied to quantify interspecific variation in head shape in a standardized way, while minimizing noise due to orientation or placement error.

As we based our analysis on landmarks placed on cephalic scales, only species from the Alethinophidia, including Colubridae, Elapidae, Lamprophiidae, Homalopsidae, Pareidae, Psammophiidae, and Pseudoxyrhophiidae were used. We excluded Scolecophidean snakes due to incompatible landmark patterns. A repeatability test was performed using ten pictures from three individuals of *Lampropeltis californiae* to explore whether the positioning of the head caused variation in landmark placement. Similarly, landmarks were placed ten times on these pictures to evaluate error in landmark placement. Variation in landmark placement due to head positioning as well as landmark placement was limited (Figure S2). Species were landmarked using the tpsDig software (Rohlf, 2015). The 2D landmark coordinates were imported into R with the *geomorph* package (Adams et al., 2025), using the function *gpagen*. Generalized Procrustes Analysis (GPA) superimposes configurations by translating, scaling, and rotating landmark coordinates to remove non-shape variation, thereby allowing comparisons of pure shape (Rohlf & Slice, 1990; Figure S1C). Species means were then computed from Procrustes-aligned coordinates and used in subsequent analyses. Semi-landmarks were slid along their tangent directions, minimizing bending energy to optimize their positions. Statistical robustness of landmarking was assessed through repeatability tests and visual inspection of variation across replicates.

### Phylogenetic signal

The phylogenetic tree was pruned to match our species in the morphometric data. To evaluate the extent to which shape variation is structured by phylogeny we used the *physignal* function in R from the *geomorph* package. The *physignal* function quantifies the strength of phylogenetic signal using a generalized K-statistic, comparing observed similarity with expectations under Brownian motion. We applied this method to test whether head shape variation was structured by shared ancestry. Significance was assessed with 1000 permutations of tip labels, generating a null distribution against which the observed signal was compared. A value close to 1 would indicate a strong phylogenetic signal, meaning closely related species exhibit more similar shapes than would be expected by random chance. On the other hand, values significantly less than 1 suggest a weaker phylogenetic signal, indicating that the variation in shape is less influenced by the evolutionary history of the species.

### Allometry

Allometry was quantified using a phylogenetic generalized least squares (PGLS) model, with shape coordinates as the dependent variable and centroid size as the predictor. This test was used to evaluate whether differences in head shape could be explained by differences in body size. Phylogenetic generalized least squares (PGLS) models fit a regression that accounts for shared ancestry by incorporating the expected covariance structure among species (Grafen, 1989). We implemented this in R using the function *procD.pgls* from *geomorph*, and statistical significance was assessed via permutation tests on residual sums of squares.

### Phylogenetic principal components analysis

A phylogenetically-informed principal component analysis (PCA) was performed using the *phytools* package (Revell, 2012) to explore shape variation in relation to habitat and diet categories. This method was applied to reduce the dimensionality of morphometric data and visualize axes of greatest variation while accounting for phylogenetic structure.

Eigenvalues were calculated to assess the variance explained by the principal components, and the first two principal components (PC1 & PC2) were visualized. The phylomorphospace was visualized by integrating the pruned phylogenetic tree and the principal component scores, with species colored based on their habitat use and diet using the *ggphylomorpho* function (Barr, 2017). Phylogenetic PCA modifies classical PCA by incorporating a phylogenetic covariance matrix, ensuring that loadings reflect evolutionary relationships. Implementation was done with the *phyl.pca* function in the *phytools* package, and significance of phylogenetic structure in ordination was assessed by comparing eigenvalues with those expected under Brownian motion simulations.

### Differences between groups

A phylogenetic multivariate analysis of variance was performed using Procrustes shape coordinates as the dependent variables to test whether habitat or diet groups differed in head shape. Phylogenetic MANOVA extends multivariate ANOVA by accounting for phylogenetic covariance, testing for mean shape differences among groups. Implementation was performed with *procD.pgls* function from *geomorph* package, which computes sums of squares and generates P-values by permutation (1000 iterations) of residuals, thereby providing a non-parametric assessment of significance.

### Analyses of covariation

A phylogenetic Two-Block PLS (Partial Least Squares) analysis was carried out using the *phylo.integration* function from the *geomorph* package, to test for covariation between head shape and diet composition. This method identifies axes of maximal covariation between two data blocks while correcting for non-independence among species due to shared ancestry (Rohlf & Corti, 2000). Implementation was performed with the *phylo.integration* function in *geomorph*, and significance was assessed through permutation tests (1000 iterations) comparing observed singular values with those from randomized datasets. We used continuous proportions of prey items (e.g., hard prey, soft prey, invertebrates, etc.), allowing for a finer representation of dietary gradients across species. This matrix was aligned with shape data from Procrustes-aligned landmarks, and phylogenetic relationships were accounted for.

### Convergence

To assess morphological convergence among species sharing similar habitats, we employed the *search.conv* function from the *RRphylo* package (Castiglione et al., 2019). This method evaluates whether species occupying the same habitat or dietary category exhibit greater phenotypic similarity than expected under a Brownian motion model of trait evolution. The analysis was conducted on a time-calibrated phylogeny and focused on the first two principal components of head shape variation (PC1 and PC2). To ensure statistical robustness, only categories represented by at least three species were retained for analysis (e.g., the soft eggs diet was not included in some analyses as they were less than 3 individuals in this group). The function quantifies convergence by calculating the mean angular distance in trait space between focal taxa and their nearest phenotypic neighbors, both in terms of trait values (ang.state) and their estimated evolutionary timing (ang.state.time). Significant convergence is inferred when observed similarity is greater than expected under neutral evolutionary expectations.

### Disparity analysis

We performed a morphological disparity analysis to assess the relationship between shape variation and habitats or diet groups while accounting for phylogenetic relationships, using the *morphol.disparity* function from the *geomorph* package. We used Procrustes-aligned shape coordinates (GPA) and tested for differences in morphological disparity across habitats and diet categories. A phylogenetically informed disparity analysis was conducted using a permutation-based resampling method (1000 iterations), enabling robust comparisons of shape variance between habitat types and diet groups.

## Results

### Phylogenetic signal

The observed value of K is low for shape (K = 0.096), indicating a weak phylogenetic signal. Furthermore, the result was not statistically significant (P = 0.91), suggesting that the morphological shape variation among species is not strongly structured by phylogenetic relationships. This implies that closely related species do not resemble each other more than expected by chance in terms of overall shape.

### Allometry

The phylogenetic generalized least squares (PGLS) analysis revealed a statistically significant relationship between shape and centroid size (Df = 1; SS = 4.12; Rsq = 0.027; *F* = 6.82; *Z* = 2.63; *P* < 0.05). Despite a relatively low proportion of variance explained, the allometric signal in head shape variation was significant.

### Variation in head shape

In our principal component analysis, 97% of the variation in head shape is explained by the first two principal components (PC1: 73.39%; PC2: 23.99%; Figure 1; Figure 2).

**Figure 1:**
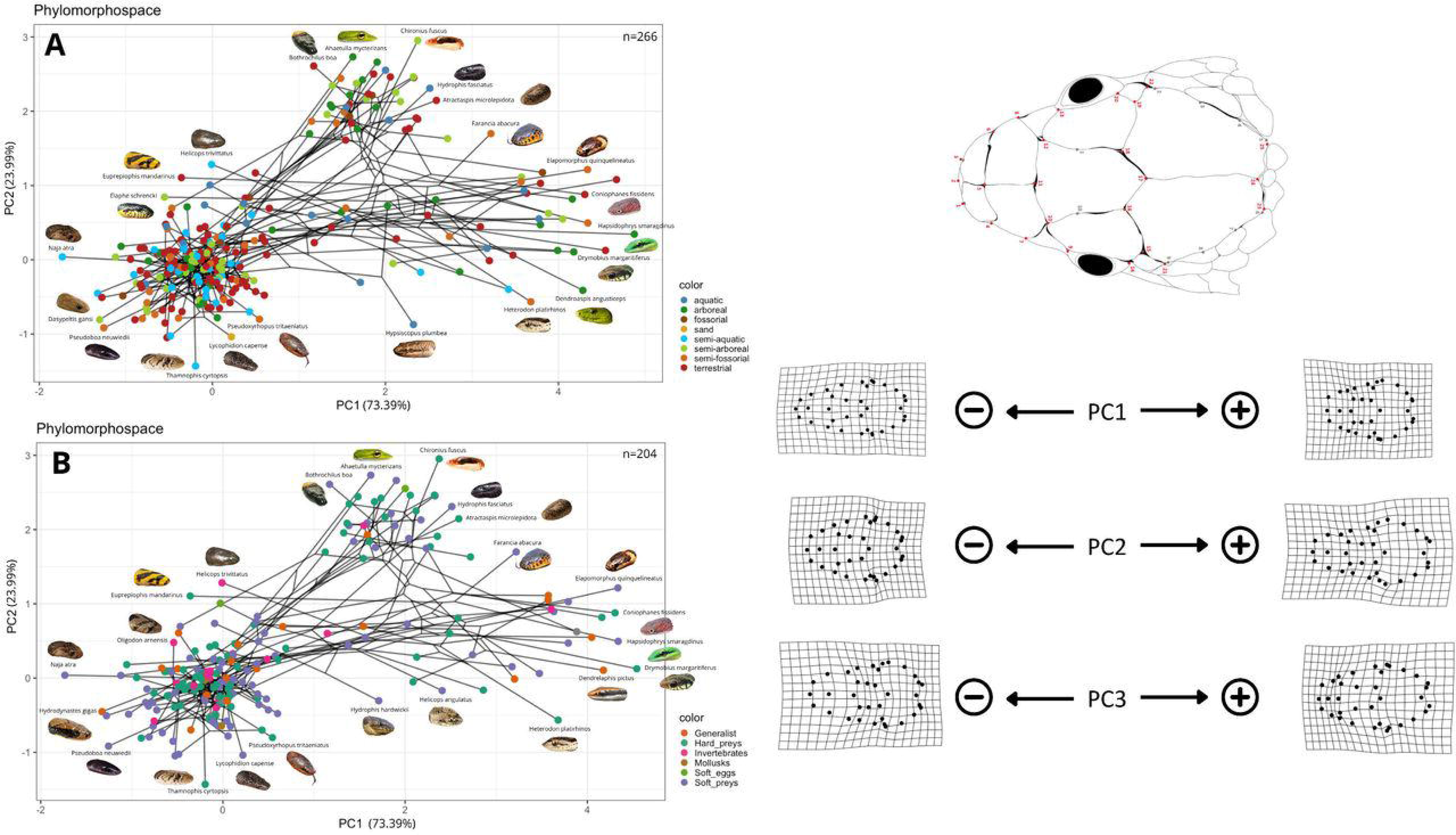
Phylomorphospace illustrating head shape variation across 281 species of snakes. The phylogeny is plotted in the morphospace described by the first two axes (PC1 vs. PC2). Nodes are colored based on species habitat classification.

**Figure 2:**
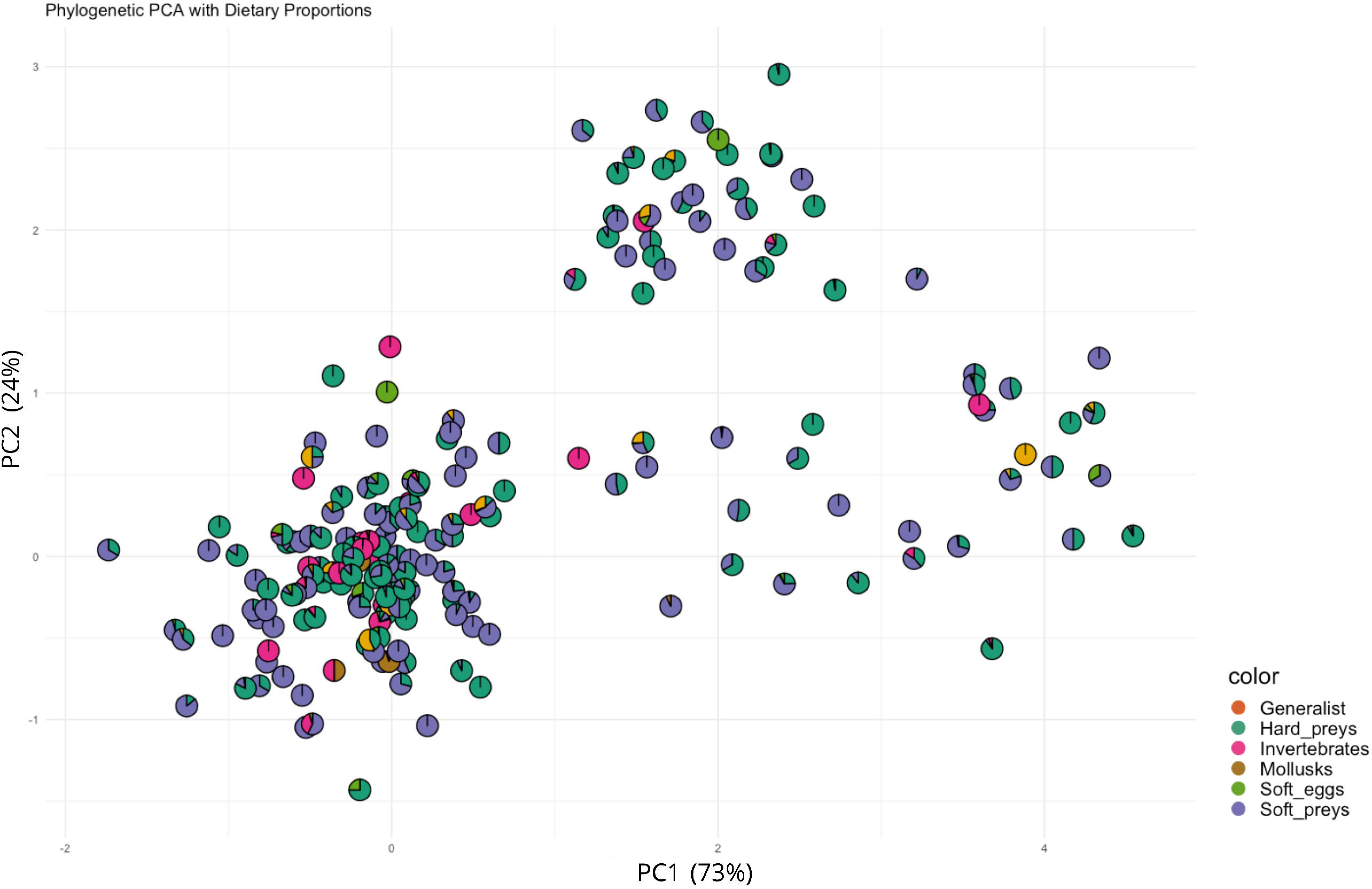
Phylomorphospace illustrating head shape variation across 197 species of snakes. The phylogeny is plotted in the morphospace described by the first two axes (PC1 vs. PC2). Nodes are colored based on species diet classification.

Although most of the variance is explained by PC1 and PC2, we also visualized the morphospaces defined by PC1 vs PC3 (Figure S3; Figure S5) and PC2 vs PC3 (Figure S4; Figure S6) to capture additional axes of shape variation and to ensure that our interpretations were not restricted to a single projection. The phylomorphospace of head landmarks reveals structuring of head shape variation among snakes. Fossorial species, characterized by low PC1 and PC2 values, exhibit elongated and broad posterior head (Figure 1). Arboreal and semi-arboreal snakes, by contrast, tend to occupy higher PC1 values and display slender, elongated heads. Aquatic and semi-aquatic species are more dispersed along PC1, with heads often appearing streamlined. Terrestrial snakes are widely spread across the morphospace and semi-fossorial and semi-arboreal species tend to fall in intermediate regions of the morphospace.

In terms of diet, species consuming soft-bodied prey (e.g., snakes, lizards) generally exhibit higher PC1 values, associated with elongated and narrower heads (Figure 2; Figure 3). Conversely, species feeding on harder prey (e.g., rodents, birds, mollusks) are found in the lower range of PC1 and often exhibit broader, more robust heads. Mollusk specialists and other durophagous feeders show this trend most strongly. PC2 variation also reflects dietary adaptations: species with lower PC2 values, such as soft-prey specialists and generalists, tend to have narrower heads, while invertebrate feeders cluster around broader and more robust head shapes. Deformation grids show that many species exhibit a combination of longer, narrower snouts with posterior heads widening—contributing to the shared V-shaped morphology in fossorial, semi-aquatic, and some hard-prey specialists (Figure 1). Overall, generalist feeders and species with diverse prey types occupy wide, overlapping regions of the morphospace, indicating that head shape in snakes is shaped by a complex interplay between habitat use and dietary specialization.

**Figure 3:**
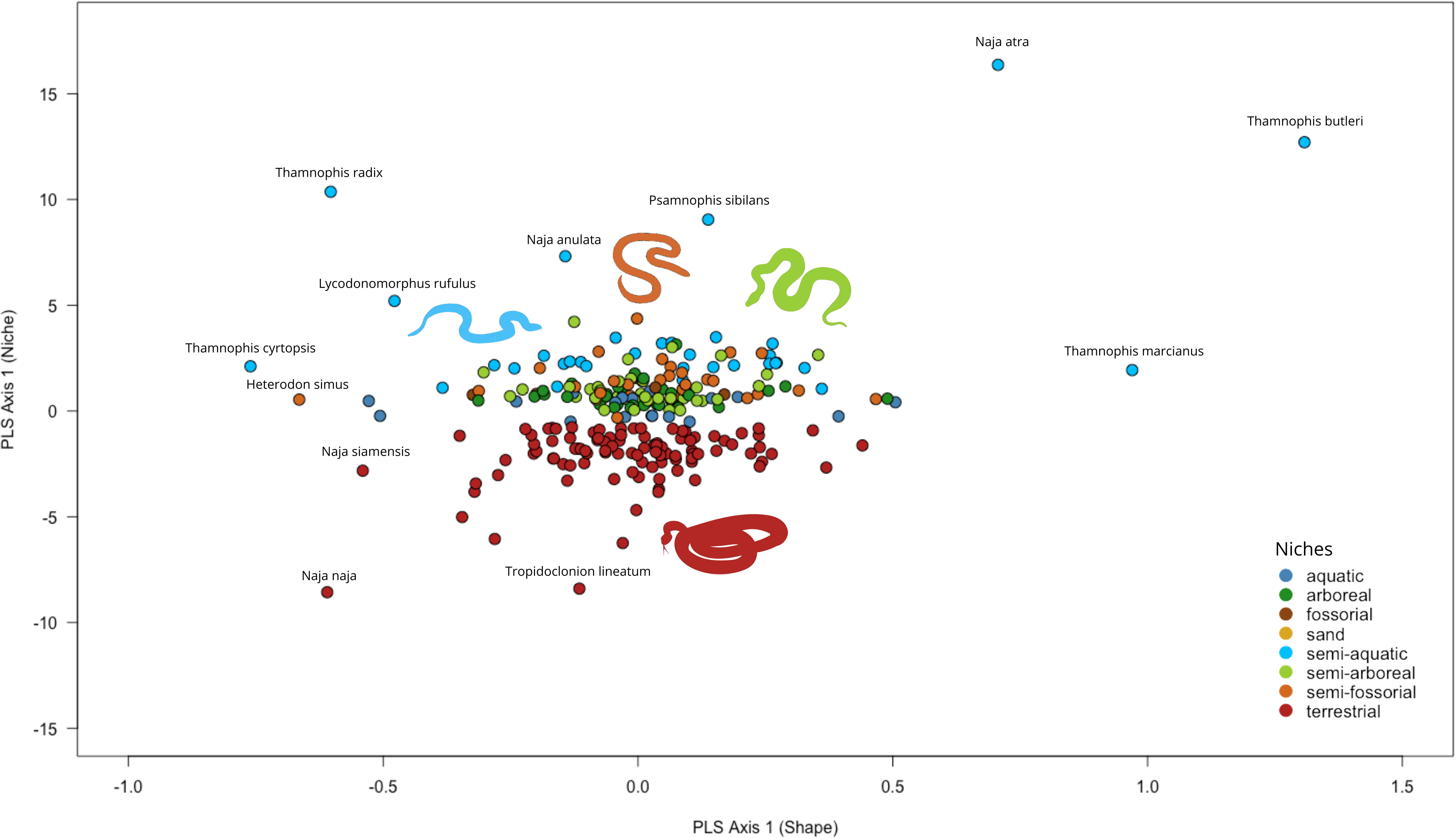
Phylogenetic PCA illustrating proportions of prey items of the 197 species of snakes included in the analyses as described by the first two axes (PC1 vs. PC2).

### Differences between habitat groups

The MANOVA indicates significant differences between habitat groups in head shape (Table 1). When adding Log10-transformed centroid size as a covariate, we found significant effects of habitat, centroid size and the interaction between the two. The interaction between habitat and centroid size was also significant. Moreover, post-hoc pairwise comparisons further revealed that fossorial snakes, as well as semi-fossorial, differ significantly from all ecological groups (Table 2). Additionally, semi-aquatic and semi-arboreal differ from terrestrial snakes in head shape.

**Table 1:** Results of the phylogenetic MANCOVA and MANOVA testing the effect of habitat use on shape with and without accounting for allometry. The table presents the degrees of freedom (Df), sum of squares (SS), mean squares (MS), proportion of explained variance (R^2^), F-statistic (F), and p-value.

| | Df | SS | MS | $R^2$ | $F$ | $P$ |
| --- | --- | --- | --- | --- | --- | --- |
| MANCOVA |  |  |  |  |  |  |
| Centroid size | 1 | 2.91 | 2.91 | 0.01 | 4.71 | $P < 0.05$ |
| Habitat | 6 | 24.94 | 4.16 | 0.13 | 6.72 | $P < 0.05$ |
| Interaction | 6 | 11.46 | 1.91 | 0.06 | 2.73 | $P < 0.05$ |
| MANOVA |  |  |  |  |  |  |
| Habitat | 6 | 24.94 | 4.16 | 0.13 | 6.33 | $P < 0.05$ |

**Table 2:** Results of the Tukey post-hoc test for pairwise comparisons of morphological disparity across habitat groups. The table presents the effect size (d), upper confidence limit at 95% (UCL 95%), Z-score (Z), and p-value for each comparison.

| Comparison | d | UCL (95%) | Z | *P* |
| --- | --- | --- | --- | --- |
| Fossorial – aquatic | 0.17 | 0.13 | 2.20 | *P* < 0.05 |
| Fossorial – arboreal | 0.17 | 0.13 | 2.17 | *P* < 0.05 |
| Fossorial – semi-aquatic | 0.18 | 0.13 | 2.28 | *P* < 0.05 |
| Fossorial – semi-arboreal | 0.17 | 0.13 | 2.25 | *P* < 0.05 |
| Fossorial – semi-fossorial | 0.20 | 0.13 | 2.40 | *P* < 0.05 |
| Fossorial – terrestrial | 0.18 | 0.13 | 2.23 | *P* < 0.05 |
| Semi-fossorial - arboreal | 0.07 | 0.06 | 2.12 | *P* < 0.05 |
| Semi-fossorial – semi-aquatic | 0.09 | 0.07 | 2.34 | *P* < 0.05 |
| Semi-fossorial – semi-arboreal | 0.06 | 0.05 | 1.94 | *P* < 0.05 |
| Semi-fossorial – terrestrial | 0.09 | 0.04 | 2.91 | *P* < 0.05 |
| Semi-aquatic – terrestrial | 0.08 | 0.06 | 2.35 | *P* < 0.05 |
| Semi-arboreal – terrestrial | 0.05 | 0.05 | 1.89 | *P* < 0.05 |

### Differences between diet groups

There was a trend toward head shape differentiation among diet groups, based on dominant dietary regimes, as revealed by the phylogenetic MANOVA (Table 3). When adding Log10-transformed centroid size as a covariate, this tendency remained. Centroid size did differ between dietary groups, while interactions were not significant (Table 3). Post-hoc pairwise comparisons indicated a significant difference in head shape between species consuming hard prey and those consuming soft prey (*P <* 0.05; Table 4). There were trends observed between hard prey and soft eggs feeders (*P* = 0.090), as well as between invertebrates and soft eggs feeders (*P* = 0.092).

**Table 3:** Results of the phylogenetic MANCOVA and MANOVA testing the effect of diet on head shape. The table presents the degrees of freedom (Df), sum of squares (SS), mean squares (MS), proportion of explained variance (R^2^), F-statistic (F), and p-value.

| | Df | SS | MS | $R^2$ | $F$ | $P$ |
| --- | --- | --- | --- | --- | --- | --- |
| MANCOVA |  |  |  |  |  |  |
| Centroid size | 1 | 5.85 | 5.85 | 0.04 | 8.59 | $P < 0.05$ |
| Diet | 5 | 6.78 | 1.36 | 0.04 | 1.99 | 0.083 |
| Interaction | 5 | 5.75 | 1.15 | 0.04 | 1.69 | 0.141 |
| MANOVA |  |  |  |  |  |  |
| Diet | 5 | 6.78 | 1.36 | 0.05 | 1.89 | 0.089 |

**Table 4:** Results of the Tukey post-hoc test for pairwise comparisons of morphological disparity across diet groups. The table presents the effect size (d), upper confidence limit at 95% (UCL 95%), Z-score (Z), and p-value (p.value) for each comparison.

| Comparison | d | UCL (95%) | Z | *P* |
| --- | --- | --- | --- | --- |
| Hard prey – Soft prey | 0.04 | 0.04 | 2.08 | $P < 0.05$ |
| Hard prey – Soft egg | 0.16 | 0.21 | 1.46 | 0.090 |
| Invertebrates – Soft eggs | 0.16 | 0.20 | 1.40 | 0.092 |

### Analyses of covariance

Our results revealed a significant covariation between head shape and habitat use (r-PLS = 0.38, Z = 2.67, *P* < 0.01), as well as between head shape and diet (r-PLS = 0.59, Z = 3.82, *P* < 0.05). This indicates that species occupying different habitat types or relying on different dietary resources differ in head shape. Specifically, the distribution of species along the first PLS axis suggests that morphological variation is aligned with ecological gradients (Figure 4). Aquatic and fossorial species appear to cluster separately from arboreal or terrestrial taxa, reflecting adaptive morphological specializations for swimming, burrowing, or climbing. In the case of diet, species that consume hard prey tend to separate from those feeding primarily on soft-bodied prey or eggs, suggesting that cranial shape variation may be driven by biomechanical demands related to prey processing (Figure 5).

**Figure 4:**
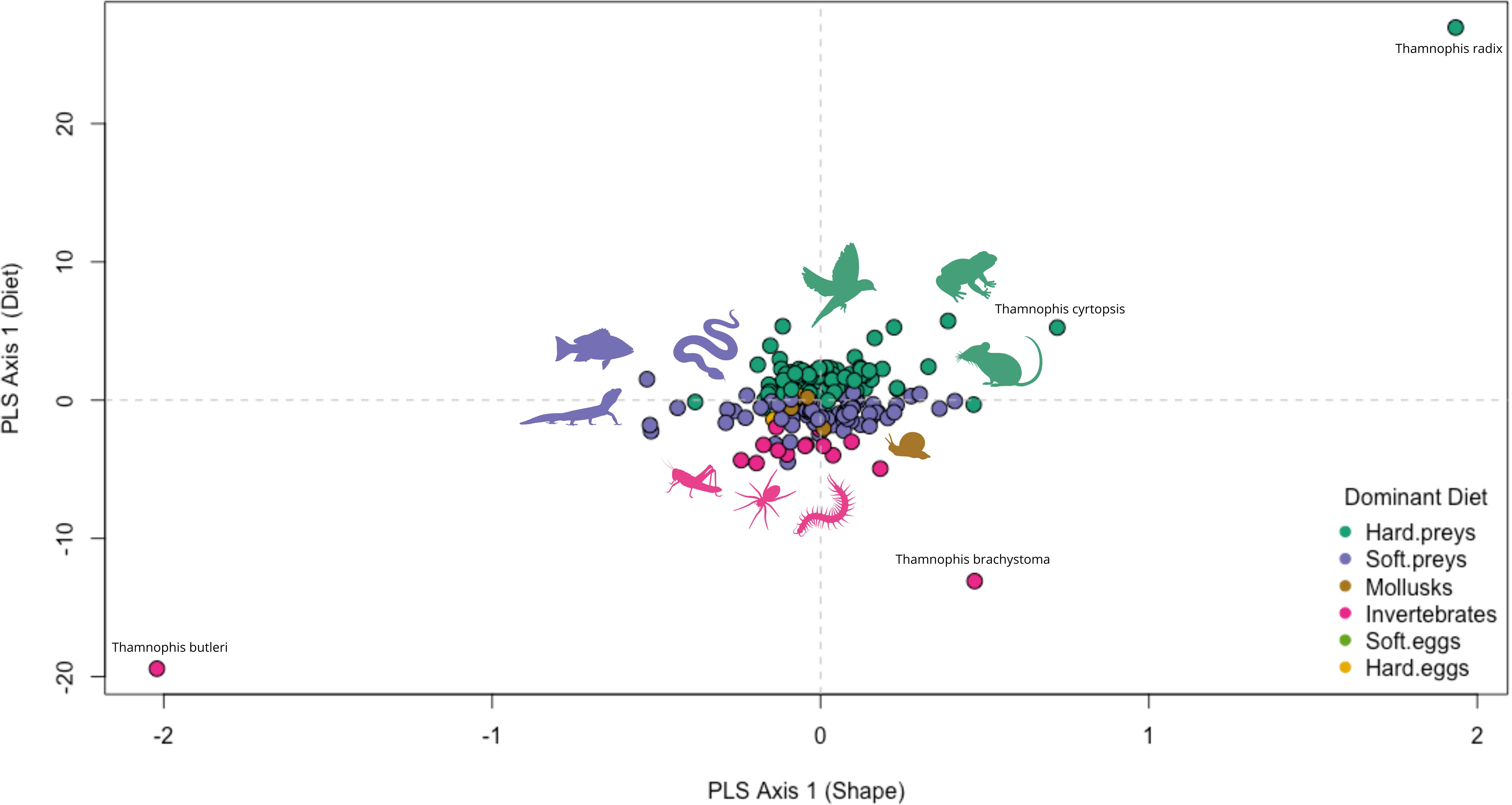
Two-block phylogenetic PLS plot illustrating covariation between head shape and ecological niches in snakes. Species are plotted along the first PLS axis of head shape (x-axis) and ecological niche (y-axis). Points are colored by habitat category, with selected species labeled.

**Figure 5:** Two-block phylogenetic PLS plot illustrating covariation between head shape and diet in snakes. Species are plotted along the first PLS axis of head shape (x-axis) and diet (y-axis). Points are colored by diet category, with selected species labeled.

### Convergence

The convergence test results revealed contrasting patterns of morphological evolution depending on whether species were grouped by habitat use or diet (Table 5). Among habitat groups, strong evidence for morphological convergence was found within aquatic and fossorial species. These two groups showed significantly lower mean angles of phenotypic distance (ang.state = 58.46 and 11.29, respectively; *P* < 0.01 for both). These results were also supported by the timing-based convergence metric, indicating that the convergent traits evolved synchronously within these clades (ang.state.time = 331.51 and 98.38, respectively; *P* < 0.01 for both groups). In contrast, no significant convergence was detected within the other habitat groups, such as terrestrial, arboreal, semi-arboreal, semi-aquatic, or semi-fossorial species (Table 5). While some of these groups (e.g., arboreal or semi-aquatic) showed moderately low angles of phenotypic distance, the results were not statistically significant.

**Table 5:** Results of the convergence test. State1 and state2 correspond to pair of ecological groups being compared. The mean angles of phenotypic distance are reported using ang.state (state) and the test of convergence with timing of diversification using ang.state.time (time), with the corresponding p-values.

|  | State | *P* | Time | *P* |
| --- | --- | --- | --- | --- |
| Habitat |  |  |  |  |
| terrestrial | 90.31 | 0.736 | 518.63 | 0.984 |
| arboreal | 85.40 | 0.087 | 478.54 | 0.271 |
| semi-arboreal | 91.41 | 0.886 | 507.65 | 0.715 |
| aquatic | 58.46 | $P < 0.01$ | 331.51 | $P < 0.01$ |
| semi-aquatic | 85.42 | 0.078 | 538.57 | 0.937 |
| fossorial | 11.29 | $P < 0.01$ | 98.38 | $P < 0.01$ |
| semi-fossorial | 90.73 | 0.563 | 525.32 | 0.850 |
| Diet |  |  |  |  |
| Hard eggs | 102.42 | 0.640 | 532.97 | 0.604 |
| Hard prey | 89.35 | 0.443 | 491.36 | 0.290 |
| Invertebrates | 87.85 | 0.304 | 508.23 | 0.621 |
| Mollusks | 53.65 | 0.165 | 448.20 | 0.372 |
| Soft prey | 88.04 | 0.155 | 501.86 | 0.616 |

No dietary group showed statistically significant morphological convergence, although mollusk specialists approached the lowest angle values (ang.state = 53.65, *P* = 0.165), hinting at possible trends that did not reach significance, potentially due to the very small sample size. All other dietary groups—including hard prey, soft prey, invertebrates, and hard egg specialists—exhibited non-significant convergence metrics. Timing-based convergence likewise yielded no significant patterns, indicating that when morphological similarities were observed, they did not evolve at the same time across lineages.

### Disparity analysis

Our findings indicate that semi-fossorial species exhibit the highest morphological disparity (0.0122; Figure 6). Aquatic species displayed the second highest score in morphological disparity (0.0116). Moderate levels of disparity were observed in terrestrial species (0.0112) and semi-arboreal species (0.0111). Conversely, fossorial species (0.0108), semi-aquatic species (0.0108) and arboreal species (0.0087) demonstrated the lowest non-zero variances. Pairwise absolute differences reveal that the greatest differences occur between aquatic and arboreal species (0.003), as well as between arboreal and semi-fossorial species (Table S5). However, pairwise absolute differences among the other habitat types remain small and statistically non-significant.

**Figure 6:** Barplot of the morphological disparity analysis across habitats and diets. The plot presents Procrustes variances for each habitat and diet, indicating the level of morphological variation within groups.

Our findings further indicated that species consuming soft prey exhibit the highest morphological disparity (0.018; Figure 6). Species consuming hard prey and mollusks displayed notable morphological disparity (0.011 and 0.0085 respectively). Moderate levels of disparity were observed in generalist species (0.0071), invertebrate feeders (0.0088) and hard-egg consumers (0.0086). Conversely, soft-egg consumers (0.0025) demonstrated the lowest non-zero variances. Pairwise absolute differences confirm that soft-prey consumers exhibit significantly higher morphological disparity compared to most other dietary categories, particularly soft-egg, mollusk, and hard-egg consumers (Table S6). Yet, pairwise absolute differences and significance levels among the other diet categories remain minimal and non-significant, except marginally for the comparison between soft-prey and soft-egg consumers.

## Discussion

Our analyses demonstrate that snake head shape variation is not primarily governed by phylogenetic relatedness or body size, but rather reflects adaptations to ecological contexts and feeding strategies. The weak and statistically non-significant phylogenetic signal (K = 0.0964) indicates that closely related species do not resemble each other in head shape more than expected by chance. This supports the idea that ecological and functional demands may override phylogenetic constraints, enabling repeated and independent morphological adaptations across lineages. Similar patterns have been observed by da Silva et al. (2018), who reported convergent heads shapes in aquatic and fossorial snakes regardless of phylogenetic affiliation, emphasizing the influence of habitat-driven functional demands. Likewise, da Silva et al. (2018) found that trophic specialization drove significant cranial divergence among Neotropical snakes with limited phylogenetic signal, suggesting that diet, like habitat, can supersede phylogenetic inertia. On the other hand, Esquerré and Keogh (2016) found a moderate phylogenetic influence on head shape in Australian pythons and boas, though ecological divergence still played a key role—highlighting potential clade-specific variation in the balance between constraint and adaptation. Although our allometric test revealed a statistically significant relationship between centroid size and shape, the effect was small (R² = 0.025). This decoupling indicates that selection acts more directly on cranial morphology than on body size (da Silva et al., 2018).

The distribution of species in morphospace reinforces this interpretation. Ecological specialization corresponds to distinct regions of cranial variation, especially among fossorial, arboreal, and semi-aquatic species. Fossorial snakes, with their V-shaped and posteriorly broadened heads, occupy a unique morphospace region—consistent with mechanical requirements for head-first burrowing and maneuvering through confined spaces (da Silva et al., 2018; Strong et al., 2020; Pandelis et al., 2023). Arboreal and semi-arboreal species exhibit slender, elongated heads, which likely facilitate navigation and prey capture in structurally complex habitats (Herrel et al., 2008; Klaczko et al., 2016). Similarly, aquatic and semi-aquatic species often show streamlined cranial shapes associated with reduced hydrodynamic drag (Segall et al., 2016, 2020; Deepak et al., 2022).

The phylogenetic MANOVA confirmed significant overall differences in head shape among habitat groups, with fossorial and semi-fossorial species showing distinct morphologies compared to most others. This pattern highlights the strong biomechanical constraints imposed by a subterranean lifestyle. Interestingly, semi-fossorial species exhibited the highest disparity among all habitat types. This elevated variation may stem from their intermediate ecological position, requiring adaptations to both surface and subsurface environments. Variability in substrate type, time spent underground, and degrees of cranial reinforcement or kinesis could contribute to this morphological diversity (Herrel et al., 2008; da Silva et al., 2018; Title et al., 2024).

Aquatic species displayed relatively high morphological disparity, likely reflecting the broad ecological diversity of aquatic environments and the wide array of prey types these snakes exploit. This is consistent with findings from Segall et al. (2016, 2020) and Deepak et al. (2022) who showed that aquatic snakes adapted to different flow regimes, prey sizes, and foraging strategies often evolve distinct cranial morphologies. Indeed, it has been shown that water acts both as a biomechanical constraint and a driver of morphological diversification (Sherratt et al., 2018). In contrast, fossorial species displayed more moderate disparity levels, which may reflect stronger functional constraints linked to a fully subterranean lifestyle. Such species often converge toward compact, reinforced head shapes adapted to head-first burrowing, thereby limiting morphological variation. In arboreal snakes, stability during strikes and secure navigation along narrow, flexible supports may favor streamlined, stable cranial morphologies (Klaczko et al., 2016; Alencar et al., 2017). Similarly, semi-aquatic species may face conflicting selective pressures between aquatic locomotion and terrestrial maneuverability, potentially resulting in conservative morphologies shaped by functional compromise (Sherratt et al., 2018; Segall et al., 2020).

Dietary specialization also contributes meaningfully to head shape variation. Species feeding on soft-bodied prey such as snakes, amphibians, or fish often display elongated, narrow heads—likely facilitating the consumption of these prey. In contrast, snakes that consume hard or bulky prey, including mammals, birds, or shelled invertebrates, possess broader, more robust heads adapted to gripping and processing resistant prey. In 2019, Perkins and Eason showed that sympatric water snake species exhibited head shapes matching their dietary preferences: narrow heads were associated with fish diets, and broad heads with amphibians. Generalists spanned a wide part of morphospace, reflecting cranial versatility and behavioral plasticity (Deepak et al., 2022; Pandelis et al., 2023). Soft-prey consumers exhibited the highest morphological disparity, likely reflecting the variety of functional solutions for handling elongate, mobile prey (Segall et al., 2020; Pandelis et al., 2023). In contrast, mollusk and soft-egg specialists clustered tightly in morphospace, suggesting convergence toward a limited set of biomechanically effective forms for durophagy (Pandelis et al., 2023).

Convergence analyses further illuminate the strong influence of habitat-driven functional constraints. Significant morphological convergence was detected among fossorial and aquatic species, likely reflecting parallel adaptations to movement through compact substrates or fluid environments (Herrel et al., 2008; Segall et al., 2016). Similarly, our detection of convergence among fossorial taxa supports the view that dealing with resistant media—whether water or soil—can lead to similar head shapes across distantly related lineages. In contrast, we did not detect significant convergence among dietary groups.

Although species consuming similar prey types often exhibited analogous head shapes, these patterns lacked statistical support for convergence. This contrasts with findings such as Perkins and Eason (2019), who documented strong diet-morphology correlations in sympatric species. The broader inconsistency may be due to variability in foraging behavior, strike mechanics, or ecological context across clades, limiting the emergence of shared morphological optima. Furthermore, lineages may face developmental or functional constraints that shape divergent cranial solutions to similar dietary challenges (Segall et al., 2020; Pandelis et al., 2023). The exclusion of underrepresented groups, such as mollusk feeders or egg specialists, from our convergence analyses may also limit our ability to detect certain trends.

Taken together, our findings reinforce the growing consensus that habitat use and dietary specialization play significant but distinct roles in shaping the evolution of head shape in snakes. Environments imposing strong biomechanical constraints, such as arboreal and fossorial habitats, clearly shape cranial form. However, diet—especially differences in prey hardness and body shape—exerts an even stronger and more consistent influence on cranial morphology across lineages, as highlighted by the PLS analysis. The pronounced covariation among head shape, habitat, and diet underscores that ecological specialization involves a dynamic interplay of selective pressures, with both habitat use and feeding ecology jointly driving morphological evolution in snakes.

While these results provide valuable insights, it is important to consider potential methodological limitations. In particular, we acknowledge the limitations of using a 2D dorsal view to represent head shape. The omission of the third dimension means that important aspects of morphology, such as underslung jaws in burrowing taxa or relatively tall heads in arboreal species, may not be adequately captured. Prior studies have shown that 2D methods can underestimate morphological variance (Buser et al., 2018) and in some cases introduce errors comparable to interspecific differences (Liu et al., 2014). The reliability of 2D analyses is particularly limited for highly three-dimensional structures. While our 2D dataset provides valuable comparative information, we note that complementary data from lateral views or full 3D reconstructions (e.g., CT scans) would enhance the robustness of functional and evolutionary inferences.

Beyond analytical methods, a number of biological and data-driven limitations should be acknowledged. First, the categorization of habitats remains somewhat coarse and potentially subjective. For example, semi-fossorial species may share more ecological traits with terrestrial than fully fossorial snakes, and semi-aquatic species vary widely in their dependence on aquatic environments (Esquerré & Keogh, 2016; Silva et al., 2017).

Incorporating quantitative ecological data—such as habitat features (e.g., branch density, soil compaction, water salinity) or substrate use (e.g., frequency of use of forest soil, mud, or tree species)—could refine these categories and better capture the ecological continuum (Manjarrez et al., 2017b; Title et al., 2024). Second, although our dietary classifications are grounded in empirical data, they could be enhanced by integrating quantitative prey traits— such as size and hardness—that more accurately represent the functional challenges faced by predators, rather than relying solely on prey shape (Vincent et al., 2009; da Silva et al., 2018). For instance, both birds and frogs may be categorized as soft prey, yet their ecologies and physical properties differ markedly, potentially requiring distinct feeding adaptations (Segall et al., 2020; Deepak et al., 2022). Understanding the functional properties of prey would thus provide more nuanced insights into head shape evolution. Furthermore, the inclusion of behavioral variables—such as foraging strategy or strike kinematics—would significantly strengthen the link between morphology and ecological function (Herrel et al., 2008; Segall et al., 2016; Moon et al., 2019). These traits are likely to interact with head shape in complex ways that are not fully captured by static ecological or dietary categories alone. Additionally, focusing on taxonomically or ecologically specialized clades (e.g., snail-eaters, egg specialists) could shed light on extreme morphological adaptations that illustrate more clearly how head shape evolves under strong selective pressure (Pandelis et al., 2023; Palci et al., 2024). Finally, subadult specimens can change our biological interpretation because of ontogenetic changes in shape where the adult form can be very different in head shape pattern compared to juveniles or even subadult individuals, due to different habitat use or diet (Patterson et al., 2022).

Our study underscores the complex and multifactorial nature of head shape evolution in snakes, driven more by ecological function than phylogenetic history or allometry. Moon et al. (2019) emphasized the remarkable adaptability of the snake feeding system, noting that head morphology reflects not only dietary and ecological demands, but also performance constraints like gape limitation, strike kinematics, and prey handling strategies. They highlight how head shape plays a central role in functional performance—such as strike accuracy, hydrodynamics, and force generation—yet also caution that our understanding remains limited by the narrow taxonomic and ecological scope of most studies. To address this, future research should aim for a more integrative approach—linking head morphology with detailed data on strike mechanics, bite force, and prey-handling behaviors across ecological contexts. For instance, striking behaviors vary between aquatic and terrestrial environments, and bite performance has only been measured in a few species, suggesting that much remains to be explored (Bartolon et al., 2013; Penning, 2017; Moon et al., 2019). Prey manipulation could have a significant effect on the overall morphology of snakes, and could be of great interest in understanding the adaptability of certain habitats to different prey items (Crotty & Jayne, 2015). Furthermore, intraspecific variation in feeding plasticity (Manjarrez et al., 2017a; Bonnet et al., 2021; Ortiz-Medina et al., 2022), as well as ontogenetic (López et al., 2013; Patterson et al., 2022) and sexual dimorphism (Borczyk et al., 2021; Shine & Goiran, 2021) in head shape and diet, could further illuminate the selective pressures shaping head form. Expanding datasets to include internal morphology, muscle architecture, and functional performance metrics would not only validate the ecological relevance of head shape variation but also test longstanding hypotheses about convergence and specialization in snake evolution. In doing so, we can build a more mechanistic understanding of how morphology translates into ecological success.

## Conclusion

Our findings reveal that snake head shape is shaped less by phylogenetic relatedness or body size and more by ecological function and dietary specialization. Morphological variation in cranial form reflects repeated, and often convergent, adaptations to environmental demands such as burrowing, climbing, or swimming, as well as to prey capture and processing. This is especially apparent in fossorial and semi-aquatic snakes, where head shapes align with biomechanical requirements like substrate penetration or hydrodynamic efficiency, and in durophagous species, which exhibit robust heads for handling hard prey.

Despite a weak phylogenetic signal and minimal allometric influence, we detected strong associations between head shape and ecological context, reinforced by disparity patterns and PLS analyses. Intriguingly, high morphological disparity within narrow ecological categories—such as fossorial and aquatic habitats—suggests the evolution of multiple functional solutions to similar environmental challenges, likely shaped by fine-scale habitat variation, prey diversity, or biomechanical constraints.

Our use of cephalic scale-based morphometrics offers a pragmatic and ecologically relevant proxy for head shape in large comparative datasets, but we recognize its limitations. Internal cranial architecture, muscle arrangement, and soft tissue all contribute to ecological performance and remain largely unexplored in this context. Moving forward, a more integrative approach—incorporating performance data (e.g., bite force, strike dynamics), behavioral traits (e.g., foraging strategy, prey handling), and finer-scale ecological metrics— will be essential for linking form to function. Investigating intraspecific variation, ontogeny, and sexual dimorphism can further illuminate how head shape evolves under diverse selective pressures. By bridging morphology with performance and behavior, future research can deepen our understanding of how ecological demands shape evolutionary trajectories, not only in snakes but across vertebrate lineages navigating similar functional landscapes.

## Author Contribution

**David Hudry:** conceptualization, investigation, writing – original draft, writing – review and editing

**Anthony Herrel:** conceptualization, investigation supervision, writing – review and editing.

## Acknowledgments

The authors would like to thank Dennis Rödder and Morris Fleck from the Museum Alexander Koenig in Bonn, and Olivier Pauwels from the Museum of Natural Sciences of Brussels for access to specimens.

## Data Availability Statement

The data that support the findings of this study are openly available on Github at https://github.com/DavidHudry/Supplementary-Geomorphometric-Data.

