## Supplementary data for "Divergent paths, convergent heads: morphological adaptation of head shape to habitat use and diet in snakes"

Figure S1 : A. Photographic setup for geometric morphometric Analysis – B. Landmark placement on a specimen (25 landmarks and 10 semi-landmarks) – C. Scatterplot of Procrustes-aligned landmark coordinates. Each point represents an individual specimen’s landmark position after GPA, accounting for sliding semi landmarks. The X and Y axes correspond to Procrustes-aligned shape coordinates, capturing morphological variation.

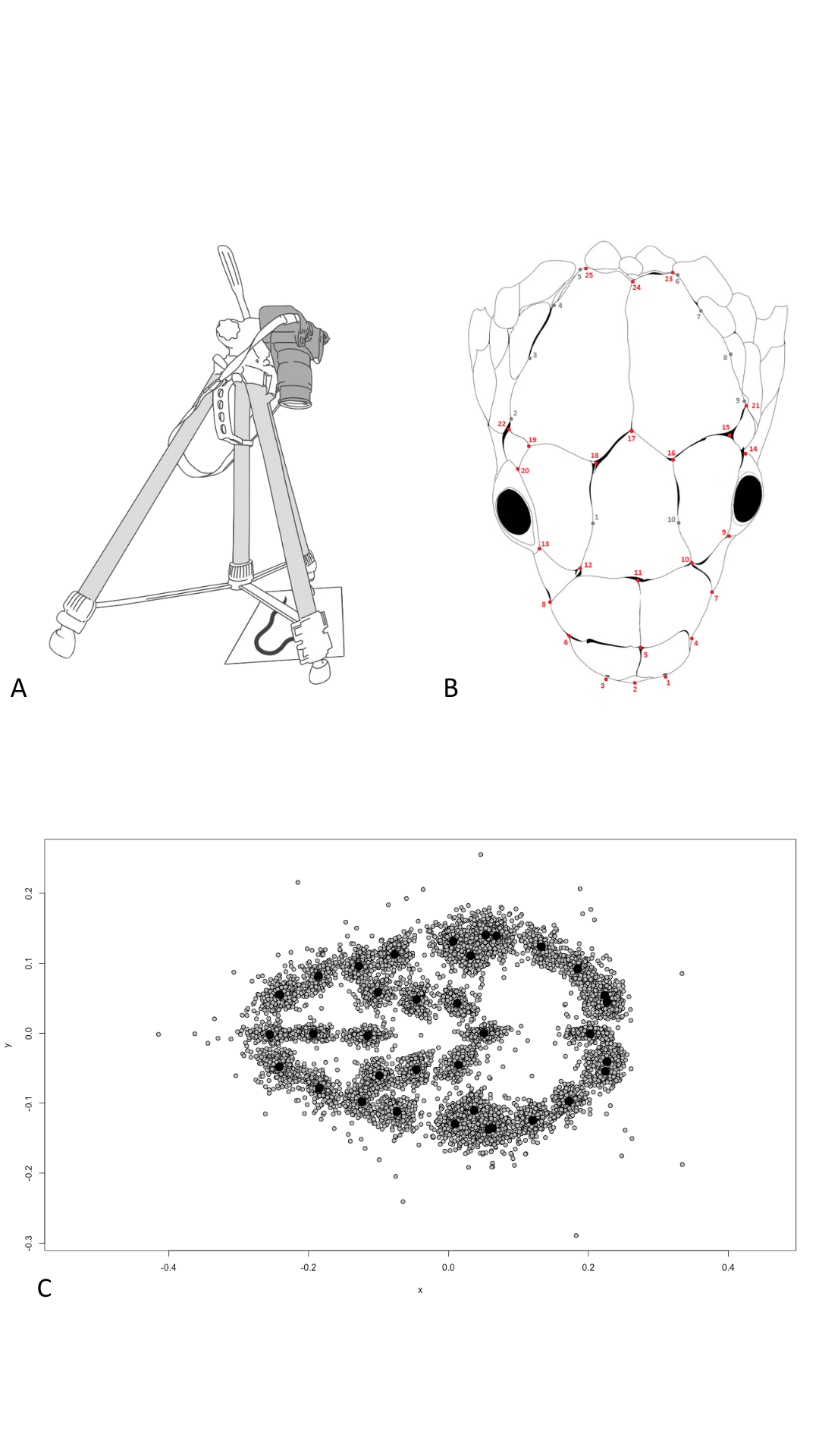

Table S1: Taxa from the dataset with non-adult individuals

| **Taxon** | **N** | **Life stage** |
| --- | --- | --- |
| *Elaphe quatuorlineata* | 2 | Young adult |
| *Pituophis melnaleucus* | 2 | Young adult |

Table S2: Definitions of habitat groups

| **Niche** | **Definition** |
| --- | --- |
| Terrestrial | Species that crawl on the ground and that rarely climb. |
| Semi-arboreal | Species that are terrestrial but with the ability to climb and move freely in vegetation. |
| Arboreal | Species that live mostly in trees and rarely come down to the ground. |
| Semi-aquatic | Species that depend on aquatic areas, mainly to feed on aquatic prey, but are generalist enough to leave these areas without difficulty. |
| Aquatic | Species that depend nearly exclusively on water. |
| Semi-fossorial | Species that live underground opportunistically or seek their food in an underground environment such as leaf litter, roots, rocks, etc... |
| Fossorial | Species that live exclusively underground. |

Table S3: Definitions of diet and composition. Diet category is defined when a prey type represents more than 50% of the total diet of the species.

| **Niche** | **Definition** |
| --- | --- |
| Generalist | Species that do not show any significant trend in prey items and cannot fit in any categories. |
| Hard eggs | Species that feed on bird eggs or reptile eggs. |
| Hard preys | Species that feed on frogs, turtles, crocodiles, mammals, or birds. |
| Invertebrates | Species that feed on annelids, ants, beetles, bugs, cockroaches, decapods, insect larvae, orthopterans, hymenopterans, arachnids, termites, or non-ant hymenopterans. |
| Mollusks | Species that feed on mollusks (mainly snails). |
| Soft eggs | Species that feed on fish eggs, or amphibian eggs & larvae. |
| Soft preys | Species that feed on salamanders, caecilians, snakes, or lizards. |

Table S4: Landmark definitions

| **Landmark** | **Definition** |
| --- | --- |
| n°1 | Anteriormost point of the snout |
| n°2 | Left lateral edge of the snout |
| n°3 | Right lateral edge of the snout |
| n°4 | Lower left edge of the first upper labial scale |
| n°5 | Lower right edge of the first upper labial scale |
| n°6 | Bottom left point of the second upper labial scale. |
| n°7 | Bottom right point of the second upper labial scale |
| n°8 | Lower left corner of the third upper labial scale |
| n°9 | Lower right corner of the third upper labial scale |
| n°10 | Midpoint between the third and fourth upper labial scales on the left side |
| n°11 | Midpoint between the third and fourth upper labial scales on the right side |
| n°12 | Upper left edge of the fourth upper labial scale |
| n°13 | Upper right edge of the fourth upper labial scale |
| n°14 | Posterior edge of the right postocular scale |
| n°15 | Posterior edge of the left postocular scale |
| n°16 | Midpoint on the upper boundary of the right ocular scale |
| n°17 | Midpoint on the upper boundary of the left ocular scale |
| n°18 | Lower left edge of the first supraocular scale |
| n°19 | Lower right edge of the first supraocular scale |
| n°20 | Upper left edge of the second supraocular scale |
| n°21 | Upper right edge of the second supraocular scale |
| n°22 | Left lateralmost point of the head, located on the parietal scales |
| n°23 | Right lateralmost point of the head, located on the parietal scales |
| n°24 | Posterior edge of the parietal scales on the left side |
| n°25 | Posterior edge of the parietal scales on the right side |

Figure S2: Repeatability test results showing ten repetitions of landmark placement for three specimens of *Lampropeltis californiae* on the PCA plot. Individuals cluster together with only minor overlap suggesting accurate landmark placement.

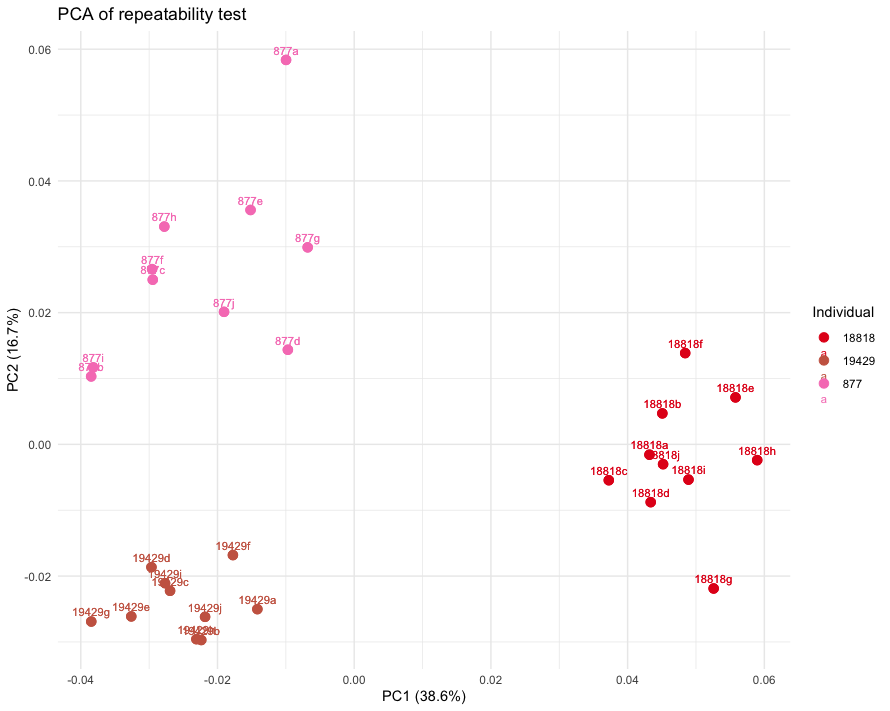

Table S5: Pairwise absolute differences in Procrustes variance (top-right side) and Pairwise p-values for Procrustes variance comparisons (bottom-left side) between habitat groups. The table displays the statistical significance of differences in Procrustes variance between pairs of habitat types.

|  | Aquatic | Arboreal | Fossorial | Semi-aquatic | Semi-arboreal | Semi-fossorial | Terrestrial |
| --- | --- | --- | --- | --- | --- | --- | --- |
| Aquatic |  | 0.003 | 0.001 | 0.001 | 0.0005 | 0.0006 | 0.0004 |
| Arboreal | 0.255 |  | 0.002 | 0.002 | 0.002 | 0.003 | 0.003 |
| Fossorial | 0.852 | 0.601 |  | 0.0001 | 0.0003 | 0.001 | 0.0004 |
| Semi-aquatic | 0.736 | 0.333 | 0.972 |  | 0.0003 | 0.001 | 0.0005 |
| Semi-arboreal | 0.839 | 0.247 | 0.964 | 0.873 |  | 0.001 | 0.0001 |
| Semi-fossorial | 0.838 | 0.106 | 0.752 | 0.535 | 0.606 |  | 0.001 |
| Terrestrial | 0.865 | 0.159 | 0.923 | 0.789 | 0.937 | 0.608 |  |

Table S6: Pairwise absolute differences in Procrustes variance (top-right side) and Pairwise p-values for Procrustes variance comparisons (bottom-left side) between diet categories. The table displays the statistical significance of differences in Procrustes variance between pairs of diet groups. Significant p-values are denoted in green.

|  | Generalist | Hard eggs | Hard prey | Invertebrates | Mollusks | Soft eggs | Soft prey |
| --- | --- | --- | --- | --- | --- | --- | --- |
| Generalist | 1.000 | 0.001 | 0.004 | 0.0002 | 0.001 | 0.005 | 0.01 |
| Hard eggs | 0.816 | 1.000 | 0.002 | 0.0002 | 0.00008 | 0.006 | 0.009 |
| Hard prey | 0.514 | 0.738 | 1.000 | 0.002 | 0.002 | 0.009 | 0.007 |
| Invertebrates | 0.734 | 0.980 | 0.644 | 1.000 | 0.0003 | 0.006 | 0.009 |
| Mollusks | 0.830 | 0.989 | 0.719 | 0.966 | 1.000 | 0.006 | 0.009 |
| Soft eggs | 0.487 | 0.396 | 0.246 | 0.328 | 0.395 | 1.000 | 0.015 |
| Soft prey | 0.067 | 0.174 | 0.216 | 0.123 | 0.172 | **0.045** | 1.000 |

Figure S3: Phylomorphospace illustrating head shape variation across 281 species of snakes. The phylogeny is plotted in the morphospace described by axes one and three. Nodes are colored based on species habitat classification.

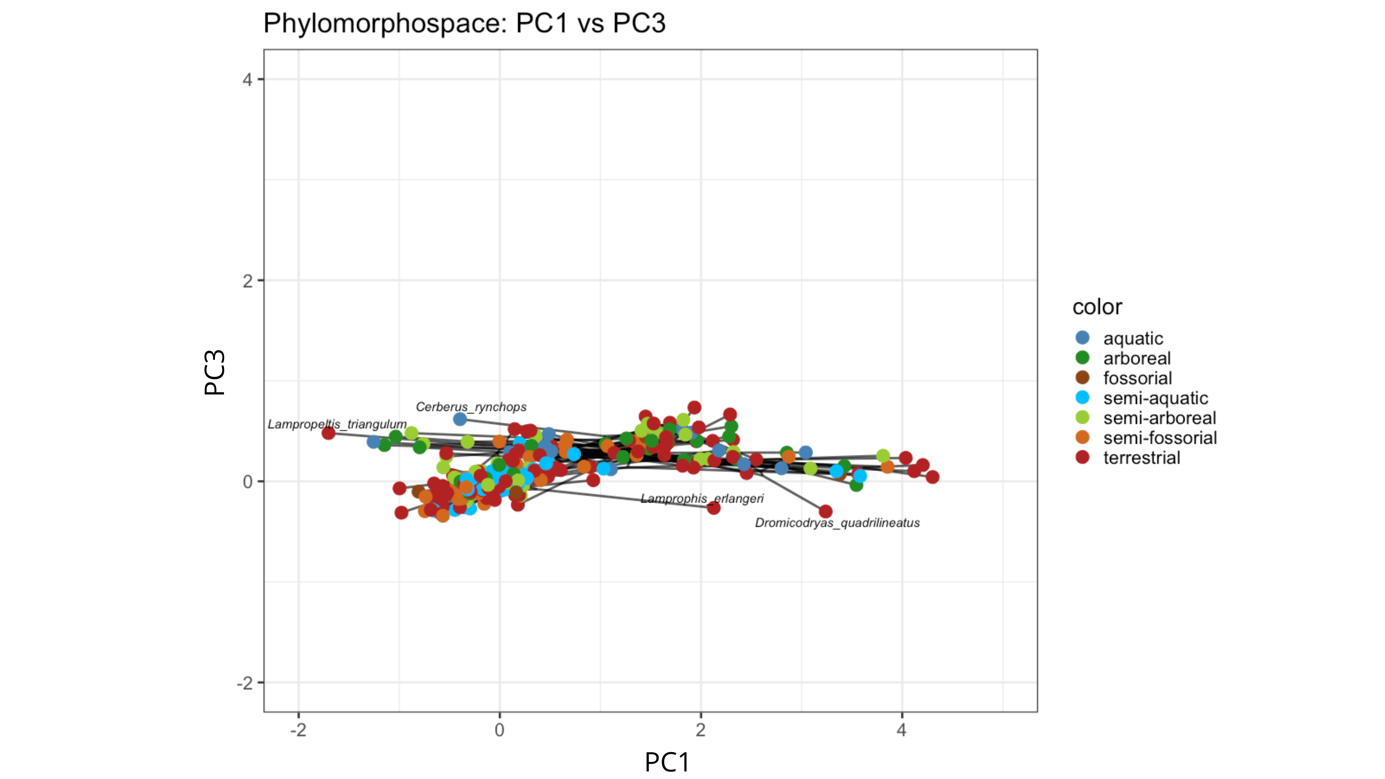

Figure S4: Phylomorphospace illustrating head shape variation across 281 species of snakes. The phylogeny is plotted in the morphospace described by axes two and three. Nodes are colored based on species habitat classification.

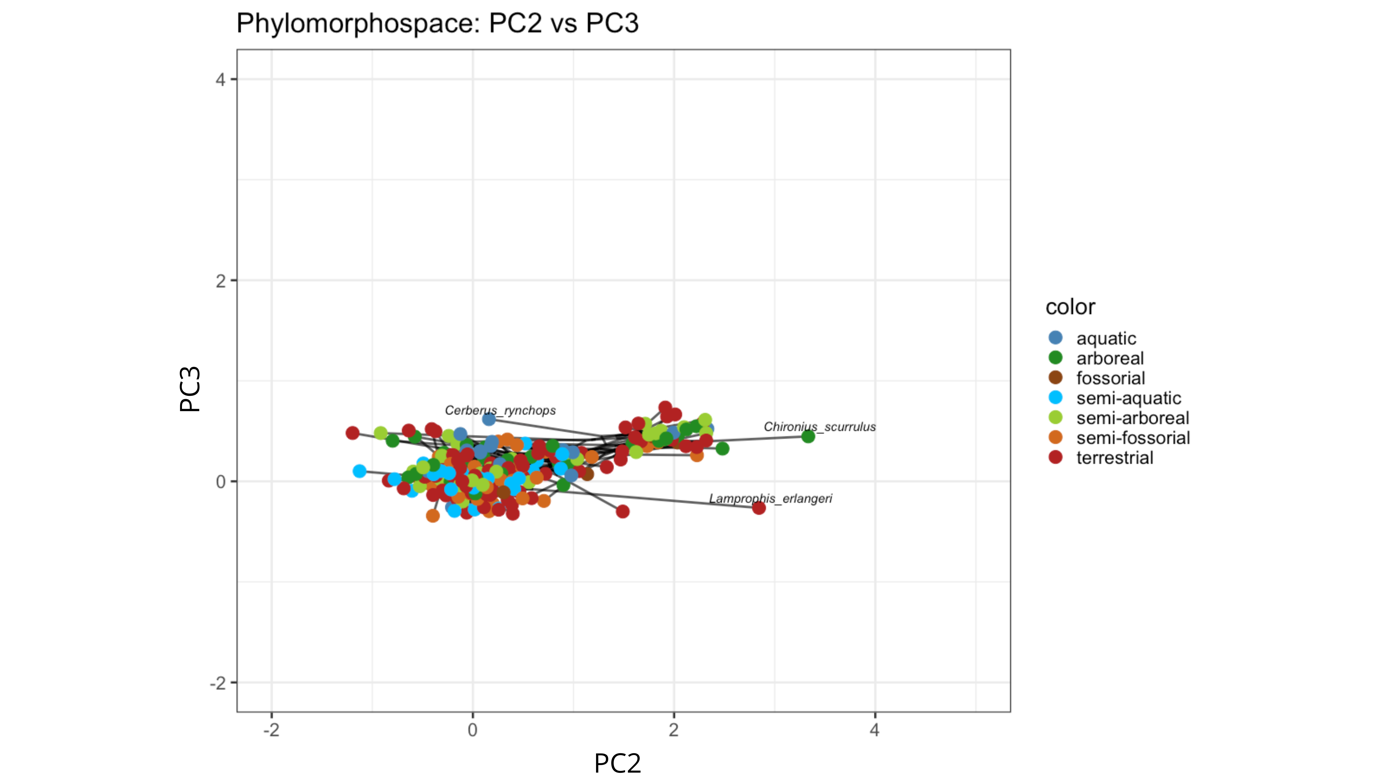

Figure S5: Phylomorphospace illustrating head shape variation across 197 species of snakes. The phylogeny is plotted in the morphospace described by axes one and three. Nodes are colored based on species diet classification.

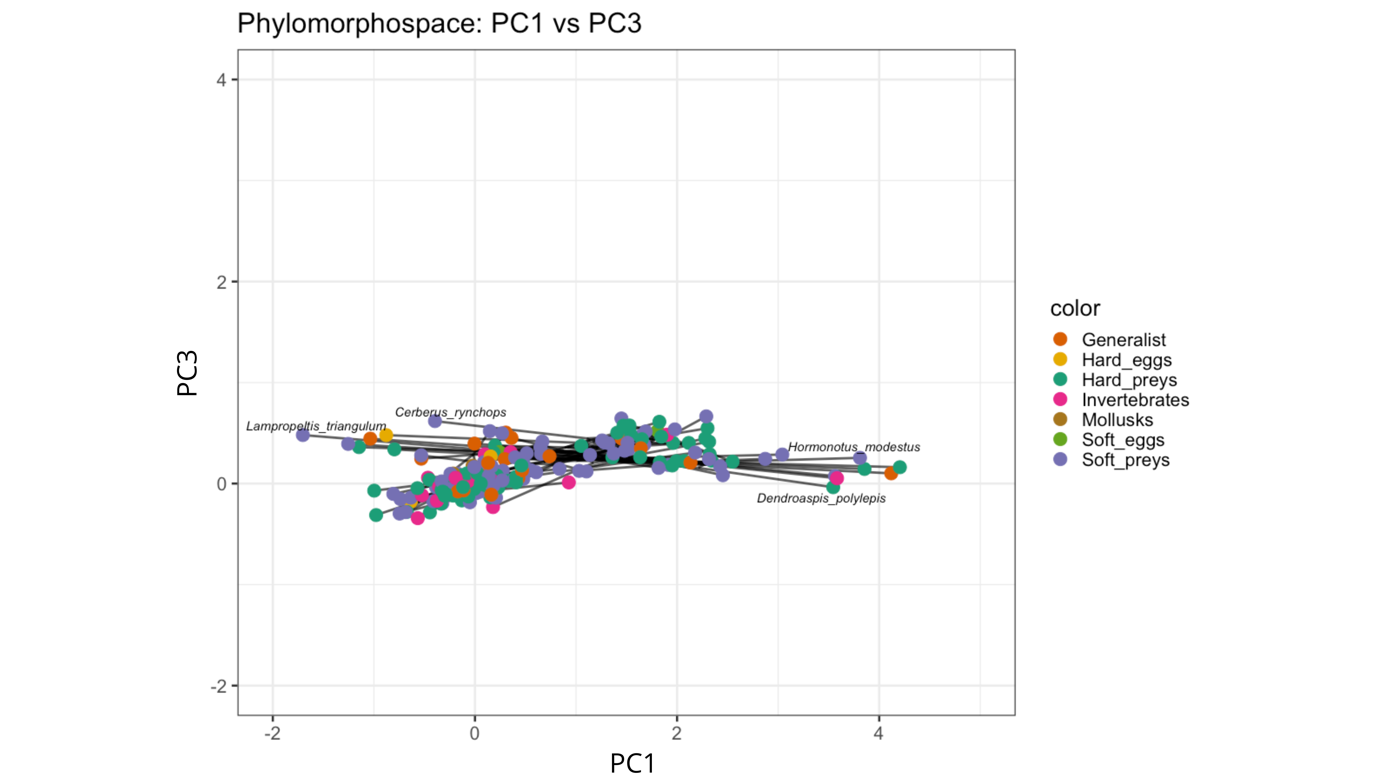

Figure S6: Phylomorphospace illustrating head shape variation across 197 species of snakes. The phylogeny is plotted in the morphospace described by axes two and three. Nodes are colored based on species diet classification.

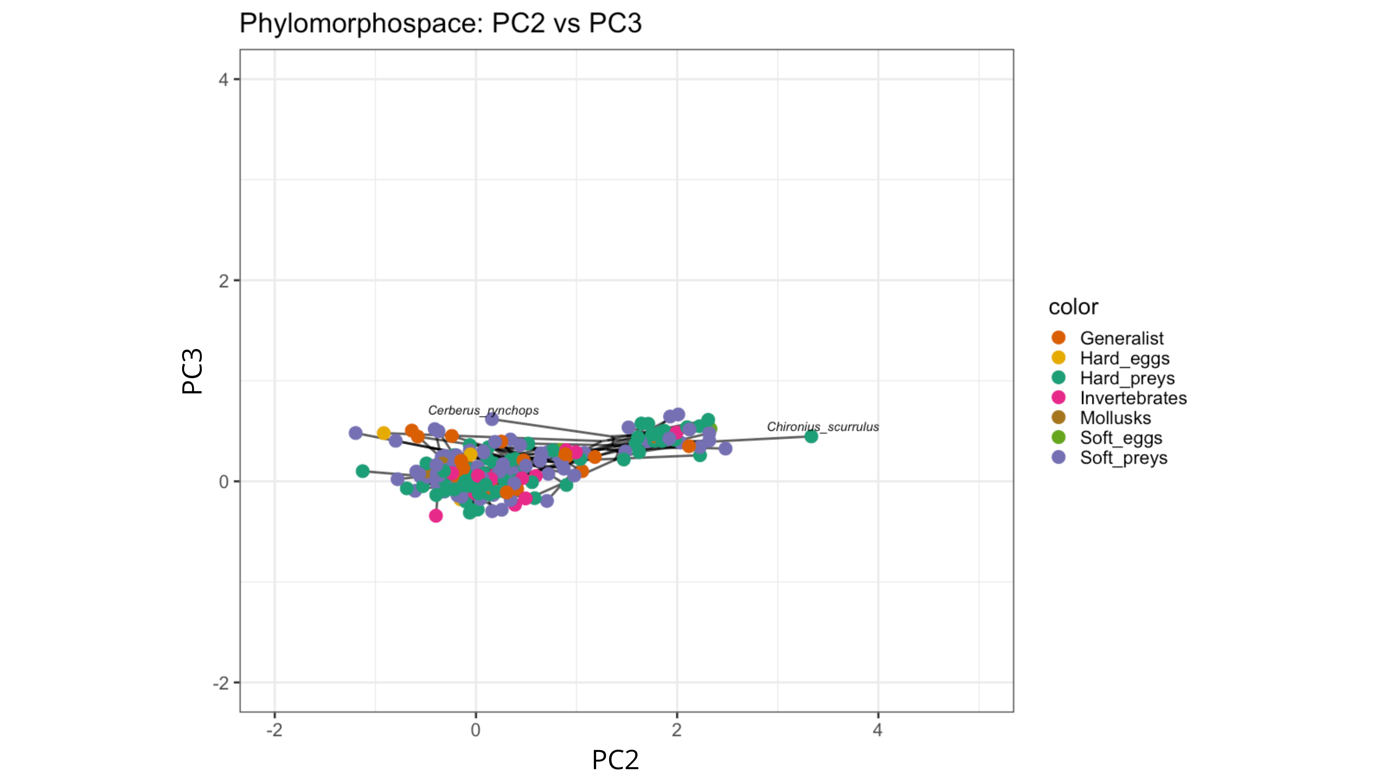
